# Enhancer/gene relationships: need for more reliable genome-wide reference sets

**DOI:** 10.1101/2022.10.12.511908

**Authors:** Tristan Hoellinger, Camille Mestre, Hugues Aschard, Wilfried Le Goff, Sylvain Foissac, Thomas Faraut, Sarah Djebali

## Abstract

Differences in cells’ functions arise from differential action of regulatory elements, in particular enhancers. Like promoters, enhancers are genomic regions bound by transcription factors (TF) that activate the expression of one or several genes by getting physically close to them in the 3D space of the nucleus. As there is increasing evidence that variants associated with common diseases are located in enhancers active in cell types relevant to these diseases, knowing the set of enhancers and more importantly the sets of genes activated by each enhancer (the so-called enhancer/gene or E/G relationships) in a cell type, will certainly help understanding these diseases.

There are three broad approaches for the genome-wide identification of E/G relationships in a cell type: (1) genetic link methods or eQTL, (2) functional link methods based on 1D functional data such as open chromatin, histone mark and gene expression and (3) spatial link methods based on 3D data such as HiC. Since (1) and (3) are costly, there has been a focus on developing functional link methods and using data from (1) and (3) to evaluate them, however there is still no consensus on the best functional link method to date.

For this reason we decided to start from the two latest benchmarks of the field, namely from the CRISPRi-FlowFISH (`CRiFF`) technique and from 3D and eQTL data in `BENGI`, and to evaluate the two methods claimed to be the best one on each of these benchmark studies, namely the `ABC model` and the `Average-Rank` method respectively, on the other method’s reference data. Not only did we manage to reproduce the results of the two benchmarks but we also saw that none of the two methods performed best on the two reference data. While `CRiFF` reference data are very reliable, it is not genome-wide and is mostly available on a cancer cell type. On the other hand `BENGI` is genome-wide but may contain many false positives. This study therefore calls for new reliable and genome-wide E/G reference data rather than new functional link E/G identification methods.

## Background

If the billions of cells of a vertebrate organism all have the same genome, their functions are as diverse as the multitude of cell types these organisms are made of. These functional differences are conveyed by the differential expression of genes across cell types, which is partly driven by differential action of their regulatory elements (promoters, enhancers, insulators, etc).

Among those regulatory elements, enhancers are particularly interesting, not only because they are predominant and cover more genomic space ([26]), but also because they appear to play important roles in human diseases ([39, 25]). Enhancers, like promoters, are DNA elements bound by transcription factors (TF). They are known to activate the expression of one or several genes by getting physically close to their promoters in the 3D space of the nucleus ([18, 31]).

There are now several catalogs of enhancers covering many different cell types, especially for human and mouse, with varying degrees of reliability. Enhancers are typically identified experimentally and *in vivo*, as for example, in the VISTA catalog ^1^, or bioinformatically, according to functional genomic data: a combination of open chromatin, histone modification and insulator data in the case of the ENCODE catalog ([23]), and Cap Analysis Gene Expression (CAGE) data in the case of the the FANTOM catalog ([1]).

The identification of all enhancers present in a given cell type is not a solved problem. The identification of the relationships between enhancers and genes, i.e. which genes are the targets of which enhancers, in a particular cell type is even more complex. However, there is increasing evidence that variants associated with common diseases are located in enhancers active in cell types relevant to these diseases ([9]). Understanding the enhancer/gene (E/G) relationships active in these particular cell types can help pinpointing important and potentially new genes associated with these diseases, and prioritizing variants in the context of genome-wide association studies ([25]). Nonetheless, this task faces important challenges because of the multivariate nature of the enhancer/gene relationship. Indeed, enhancers can (1) be far away from the genes they activate (up to several Mbp), (2) act either upstream or downstream from the activated genes and (3) activate several genes and need to be associated to other enhancers to activate a given gene ([18, 31]).

There are three broad approaches that are currently used for the genome-wide identification of E/G relationships in a given cell type (Figure 1): (1) genetic link methods that identify eQTL genetic variants, potentially located in regulatory elements such as enhancers, using expression data (microarray, RNA-seq) applied to a given cell type ([3, 17]), (2) functional link methods that directly identify E/G using genome-wide functional genomic 1D data (open chromatin, histone mark, TF, expression) in one or several cell types (see next section), and (3) spatial link (3D) methods that predict E/G using a combination of genome-wide 1D and 3D data (promoter capture HiC, ChiA-PET, etc) in a given cell type, considering The identification of all enhancers present in a given cell type is not a solved problem. The identification of the relationships between enhancers and genes, i.e. which genes are the targets of which enhancers, in a particular cell type is even more complex. However, there is increasing evidence that variants associated with common diseases are located in enhancers active in cell types relevant to these diseases ([9]). Understanding the enhancer/gene (E/G) relationships active in these particular cell types can help pinpointing important and potentially new genes associated with these diseases, and prioritizing variants in the context of genome-wide association studies that true E/G relationships of a given cell type are subsets of pairs of genomic regions predicted as close in the 3D space of the nucleus ([16, 34]).

**Figure 1:**
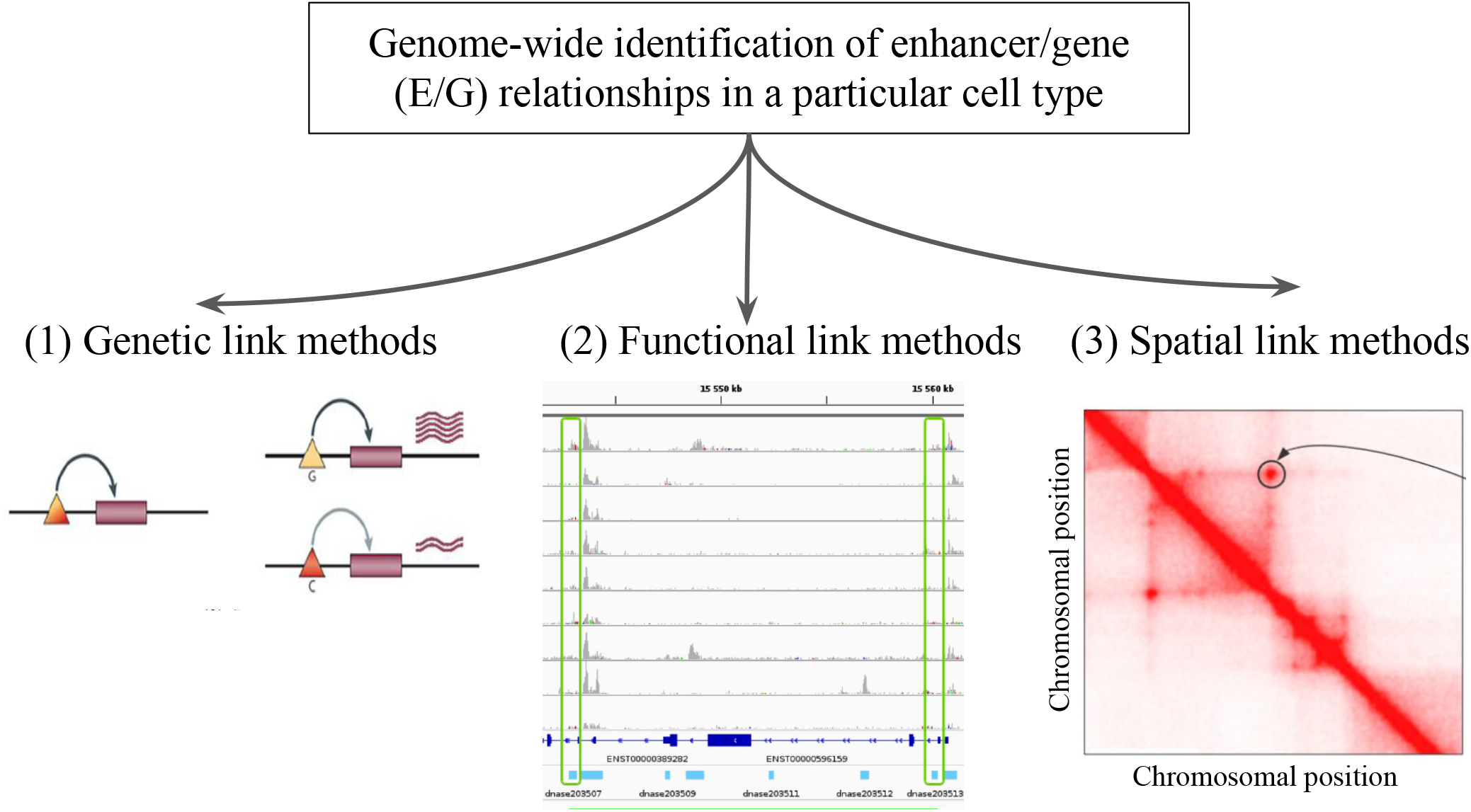
The genome-wide identification of enhancer/gene (E/G) relationships in a particular cell type. Illustration of the three broad approaches that have been described in the literature: (1) genetic link methods, (2) functional link methods and (3) spatial link methods. In panel (1) the triangles and rectangles represent genetic variants and genes, respectively. When the variant is G the gene is highly expressed, and when it is C the gene is lowly expressed. This variant is said to be an eQTL of the gene, and if located in an enhancer the relationship between the variant and the gene becomes an E/G. Panel (2) illustrates a typical heuristic functional link method, which correlates chromatin accessibility in promoters and enhancers across several cell types and is described in more details in Figure 2 below. Panel (3) represents a squared heatmap where both the horizontal and the vertical axes represent the same portion of the genome divided into equal size bins. The darker the red in the cell, the closer the two regions in the 3D space of the nucleus according to HiC data. Apart from the diagonal, some points far from the diagonal indicate relationships that could be E/G if one of the bin lies in an enhancer and the other one lies at the transcription start site (TSS) of a gene.

Because genetic (1) and spatial link (3) methods are very costly and the generation of 3D data in spatial link methods requires a specific expertise, functional link methods (2) have become the most widely used approach to identify E/G relationship. This is confirmed by the plethora of functional link methods that have been developed since 2011 (see below). On the other hand, the data underlying methods of types (1) and (3) are commonly considered as a reference to assess the reliability of methods of type (see below).

## Results

### Functional link methods for the genome-wide identification of enhancer/gene (E/G) relationships in a particular cell type

Functional link methods can be divided into two broad categories, non supervised / heuristic methods and supervised machine learning methods. While the former generally use few types of functional genomic data in a large number of cell types, the latter use many types of functional genomic data in a reduced number of cell types. Broadly speaking, non supervised methods use correlations between functional genomic signals present at enhancers and promoters across many cell types. Distance between promoters and enhancers as well as correlation threshold are determined heuristically and the evaluation of the accuracy of the method is done *a posteriori* using external reference data (most often 3D or genetic) ([10, 33, 35, 32, 1, 8, 38, 12, 22]) (Figure 2). The second category of methods use machine learning approaches such as random forests or neural networks. They first train a model to discriminate true vs. false E/G based on distinctive features from the 1D data using a reference dataset of known E/G (most often determined using a combination of 1D data for enhancer and promoter identification and 3D or genetic data for the relationship identification) and a dataset of unsupported E/G as a negative control. When provided with a new relationship associated to 1D data attributes, the model is used to decide whether the relationship is more likely to be a true E/G than a false E/G ([29, 2, 14, 30, 36, 6, 37, 13, 20, 4, 15, 11]) (Figure 3).

**Figure 2:**
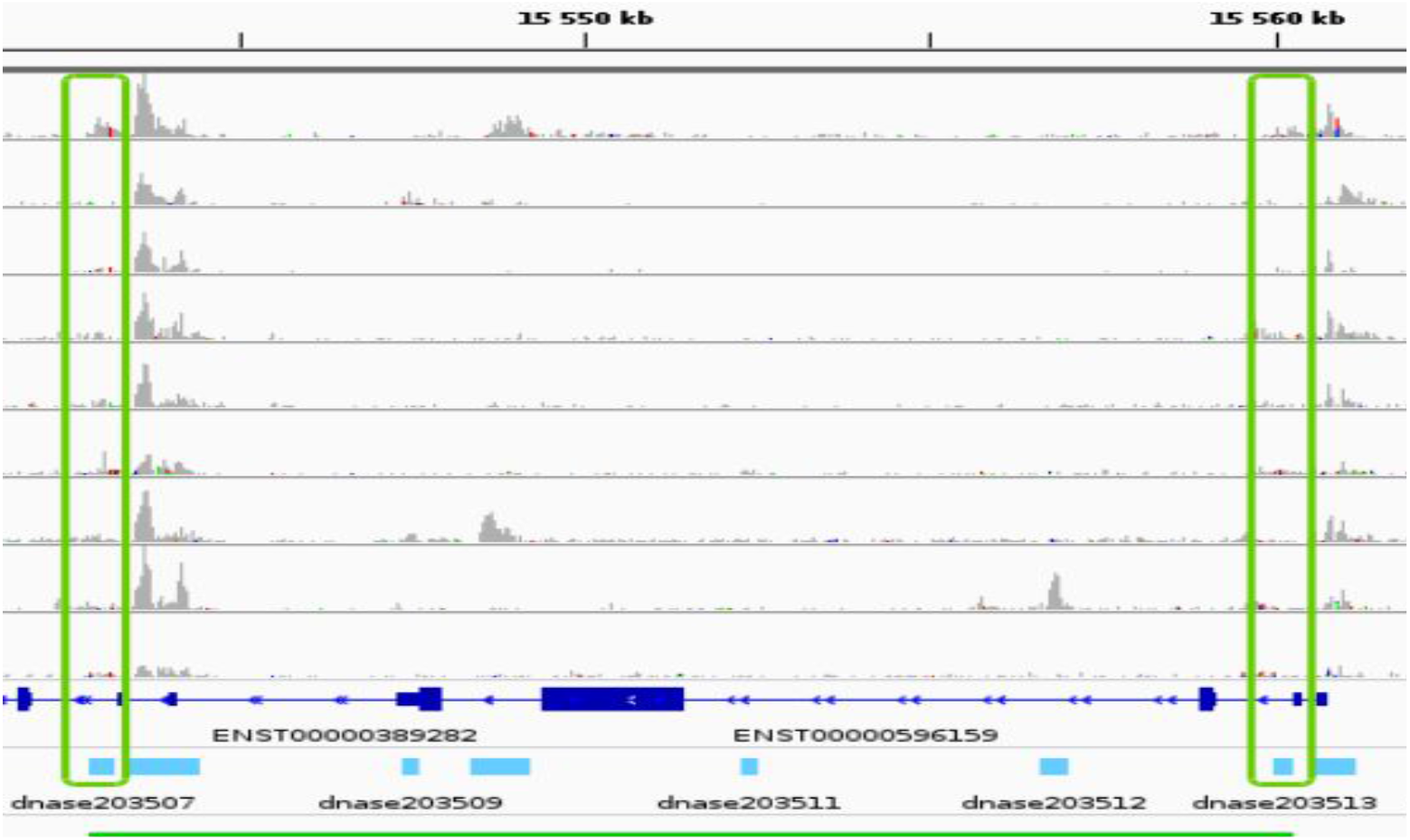
Example of a non supervised/heuristic method for the identification of E/G in a cell type. A common approach to identify candidate E/G is to investigate the correlation between chromatin accessibility signal at two regions across several cell types. The plot represents a portion of the human genome (from position 15,540 kb to position 15,560 kb of chromosome 19 on the hg19 human genome assembly) in IGV. The horizontal tracks represent ENCODE DNAse-seq signal in 10 different cell types (stomach, HepG2, K562, thymus, adrenal gland, small intestine, GM12878, IMR-90, heart and H1-hESC), followed by gene annotation (Gencode v19) in dark blue and by DNAse-seq consensus peaks from the 10 cell types in light blue. The vertical green rectangles highlight two consensus peaks (and their signal) that have a high (more than 0.7) Pearson correlation between the log10 of their normalized openness across the 10 cell types (see Material and Methods). Because the consensus peak is at the transcription start site (TSS) of a gene, it is typically interpreted as an E/G.

**Figure 3:**
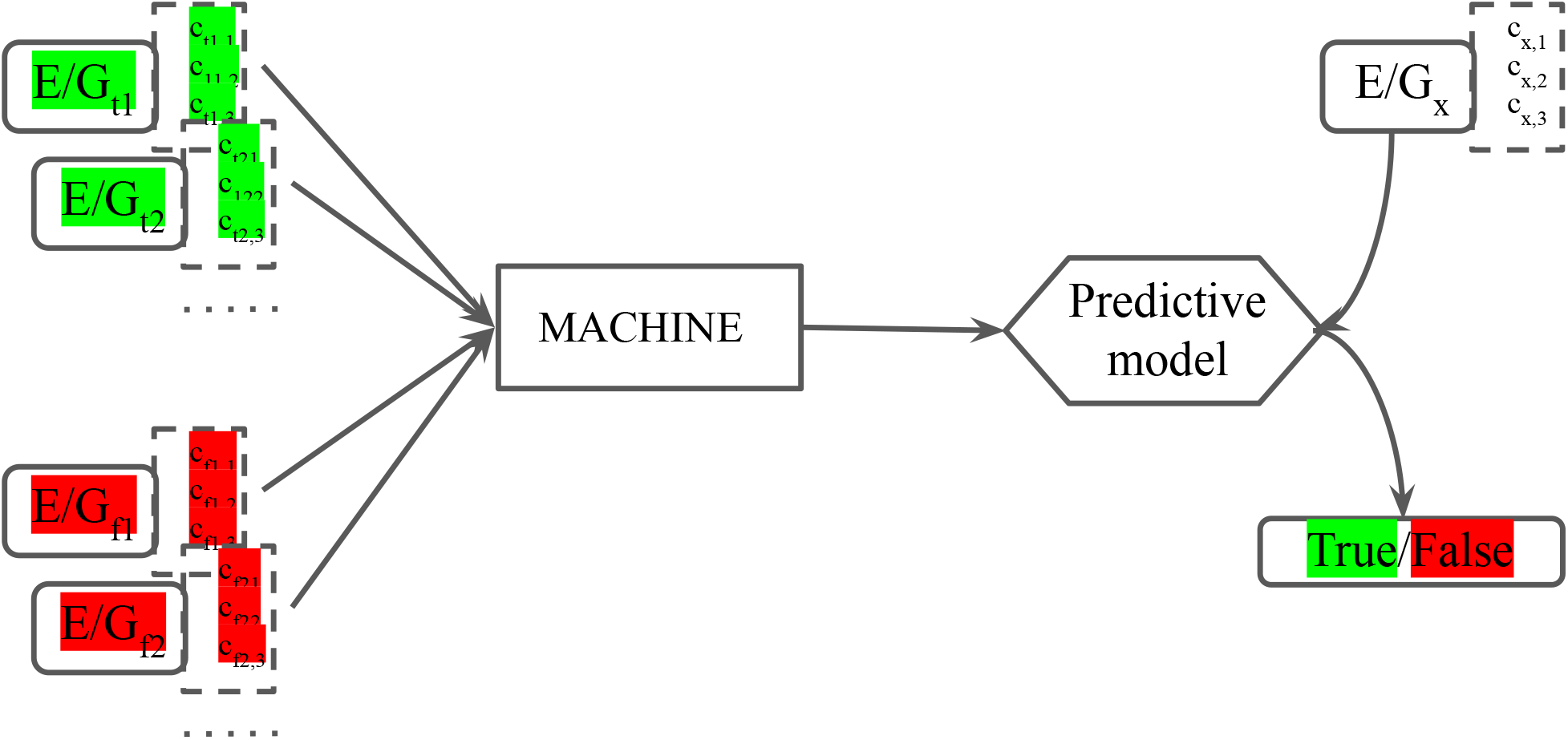
Archetype of a supervised machine learning method for the identification of E/G in a cell type. Supervised method for the identification of E/G in a cell type works in two steps, a training step (left) and a predictive step (right). In the training step, the machine is provided with true E/G in green and false E/G in red. Each E/G is provided along with a list of features, typically from functional genomic 1D data such as the accessibility of the chromatin at the enhancer and at the promoter of the gene, the gene expression level, etc, so that the machine can build a discriminative model between true and false E/G. In the predictive step, the discriminative model is fed with a new E/G and its attributes, and predicts whether this new E/G is real (true).

### Evaluating the most recent functional link methods

Two recent studies evaluated functional link methods ([12, 22]). However they do not evaluate the same methods and do not rely on the same reference data for their evaluation. Each study proposes a new method that is reported as the most accurate on the reference set used ([5]). In order to compare the performances of existing methods in an unbiased way, we therefore decided to cross-evaluate these two methods on the two reference sets provided by the two aforementioned studies ([12, 22]). First it has to be noted that those two methods are both unsupervised/heuristic: the first one, proposed by Fulco et al, is called the `Activity-By-Contact (ABC)` model ([12]), and the second one, proposed by Moore et al, is called the `Average-Rank` method ([22]).

The `ABC model` defines the score of a potential E/G in a cell type as the product of the activity of the potential enhancer E in this cell type, and the contact between E and gene G, divided by the sum of the same products but across all potential enhancers in a 5Mb region from G. The potential enhancers of a cell type are the chromatin accessible peaks in this cell type, and the aim of the 1st step of the method is to find them all. The activity (A) of E is then further computed as the geometric mean of the read counts of chromatin accessibility (usually assessed using DNAse-seq or ATAC-seq) and H3K27ac ChIP-seq at E. The contact (C) between E and G is then computed either as the Knight-Ruiz (KR) matrix-balancing normalized Hi-C contact frequency between E and the promoter of gene G, if cell type specific Hi-C data is available, or simply as the inverse of the distance (fractal globule model) between E and G ([12]). In order to predict E/G only for expressed genes, the `ABC model` can either take cell type specific gene expression data, or consider as a proxy of gene expression the activity of its promoter as defined above using chromatin accessibility and H3K27ac ChIP-seq data.

The `Average-Rank` method defines the score of a potential E/G as the inverse of the average of the ranks provided by the `Sheffield` and the `distance` methods ([22]). The `Sheffield` method was introduced in 2012 and defines the score of a potential E/G as the Pearson correlation between the chromatin accessibility at E (assessed by DNAse-seq) and the expression of G across 112 cell types ([32]). The `distance` method’s score of a potential E/G is the inverse of the distance between E and G. Here potential enhancers are all distal enhancer elements (dELS) of the ENCODE registry of candidate cis-regulatory elements (`cCREs`) ([23]).

The two evaluations mentioned above also proposed their own reference/evaluation datasets.

Fulco et al’s reference dataset is based on the output of a new genetic screening technique developed by the authors, and specifically designed to predict E/G in a cell type for a small number of genes: the `CRISPRi-FlowFISH (CRiFF)` technique ([12]), that they apply to the K562 cancer cell line. As stated by the authors, this technique perturbs “hundreds of noncoding elements in parallel and quantify their effects on the expression of an RNA of interest, combining CRISPR interference, RNA fluorescence in situ hybridization (FISH) and flow cytometry”. In this approach, they “deliver KRAB-dCas9 to many candidate regulatory elements in a population of cells by using a library of guide RNAs”. The results of this technique are then subjected to a statistical framework to determine the sets of E/G that are active and inactive in the cell type. The technique was then applied to 30 genes in five genomic regions (spanning 1.1–4.0 Mb) for which they tested all DNase I hypersensitive (DHS) elements in K562 cells within 450 kb. This approach yielded 109 positive and 3,754 negative E/G, which are then considered as the reference set for the evaluation of numerous methods of the field including the `ABC model` and the `distance` ([12], Table 1).

**Table 1:**
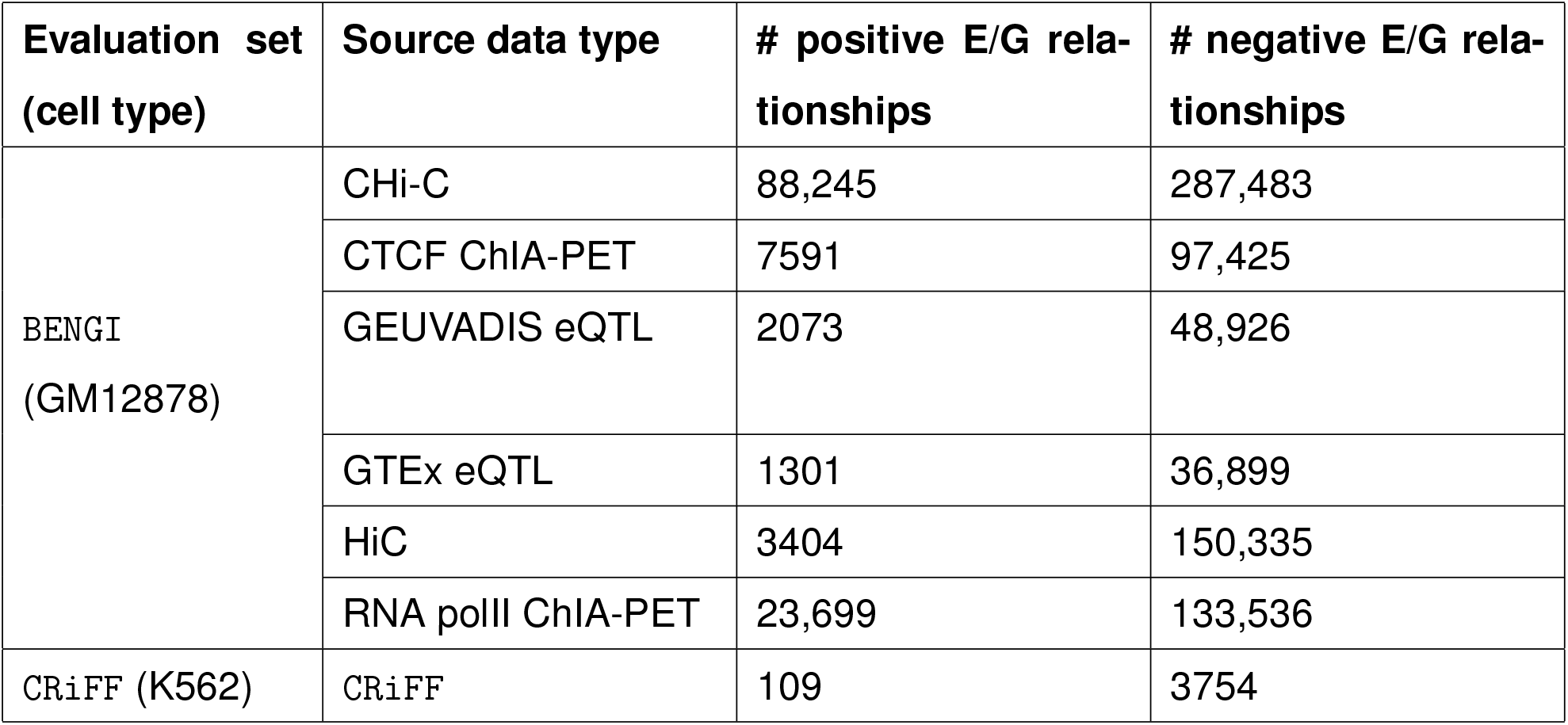
Number of positive and negative E/G relationships for each of the two evaluation sets considered.

Moore et al’s reference dataset is called `BENGI` (Benchmark of candidate Enhancer-Gene Interactions) and is made of 6 sets of E/G active and inactive in the lymphoblastoid cell line GM12878. These 6 sets of active E/G in GM12878 result of the processing of four types of 3D data, Hi-C ([27]) and promoter capture Hi-C ([21]) data and ChiA-PET of polymerase II and CTCF ([34]) data, and of eQTL data from 2 different studies, GEUVADIS ([19]) and GTEx ([7]). The negative sets are built by taking for each enhancer of a positive set, all the genes not connected to it in the positive set and lying within the 95 percentile of the positive set distances from it. The number of positive and negative E/G obtained are indicated in Table 1. Since 3D and eQTL data are not specifically generated to identify E/G relationships, the `BENGI` reference sets are expected to be overall less reliable than the `CRiFF` reference set. However the fact that `BENGI` provides genome-wide information is an advantage over `CRiFF`.

Given all these data we proceeded to the evaluation of the `ABC model`, the `Average-Rank` and the `distance` methods on both the `BENGI` sets and the `CRiFF` set, using the code provided by the authors of the two papers (with some adjustments), and obtained the results presented on Figures 4 and 5, respectively (see Material and Methods). It has to be noted that only Fulco et al provided a code dedicated to their proposed method ^2^, Moore et al only provided code dedicated to the evaluation of their proposed method on any evaluation set ^3^.

**Figure 4:**
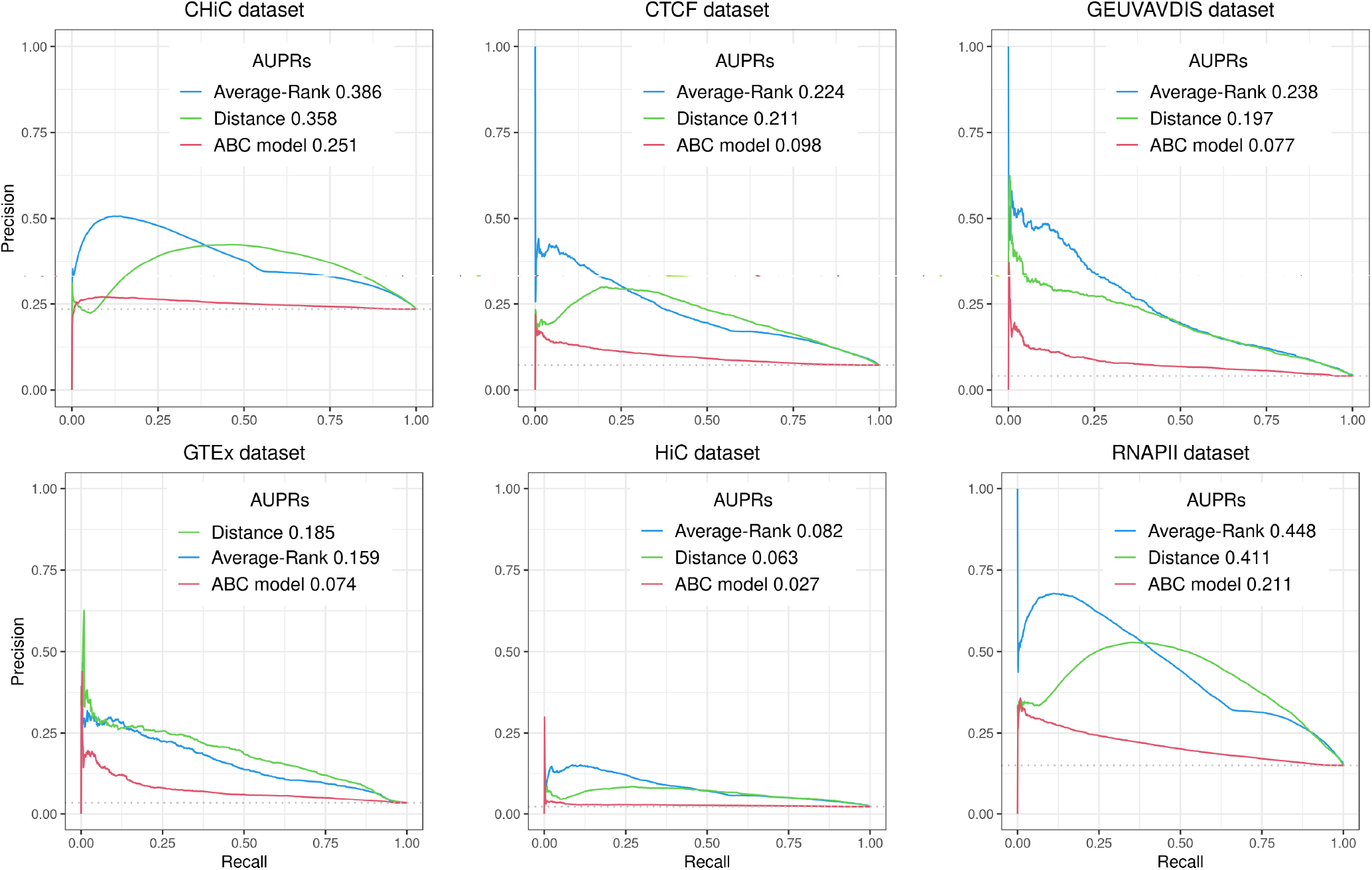
Precision-Recall curves for the `Average-Rank`, the `distance` and the `ABC model` methods on the six datasets of the `BENGI` evaluation set (all pairs, natural ratio). AUPRs (Area Under the Precision-Recall curve) of each method are also indicated on the top right end corner of each plot.

**Figure 5:**
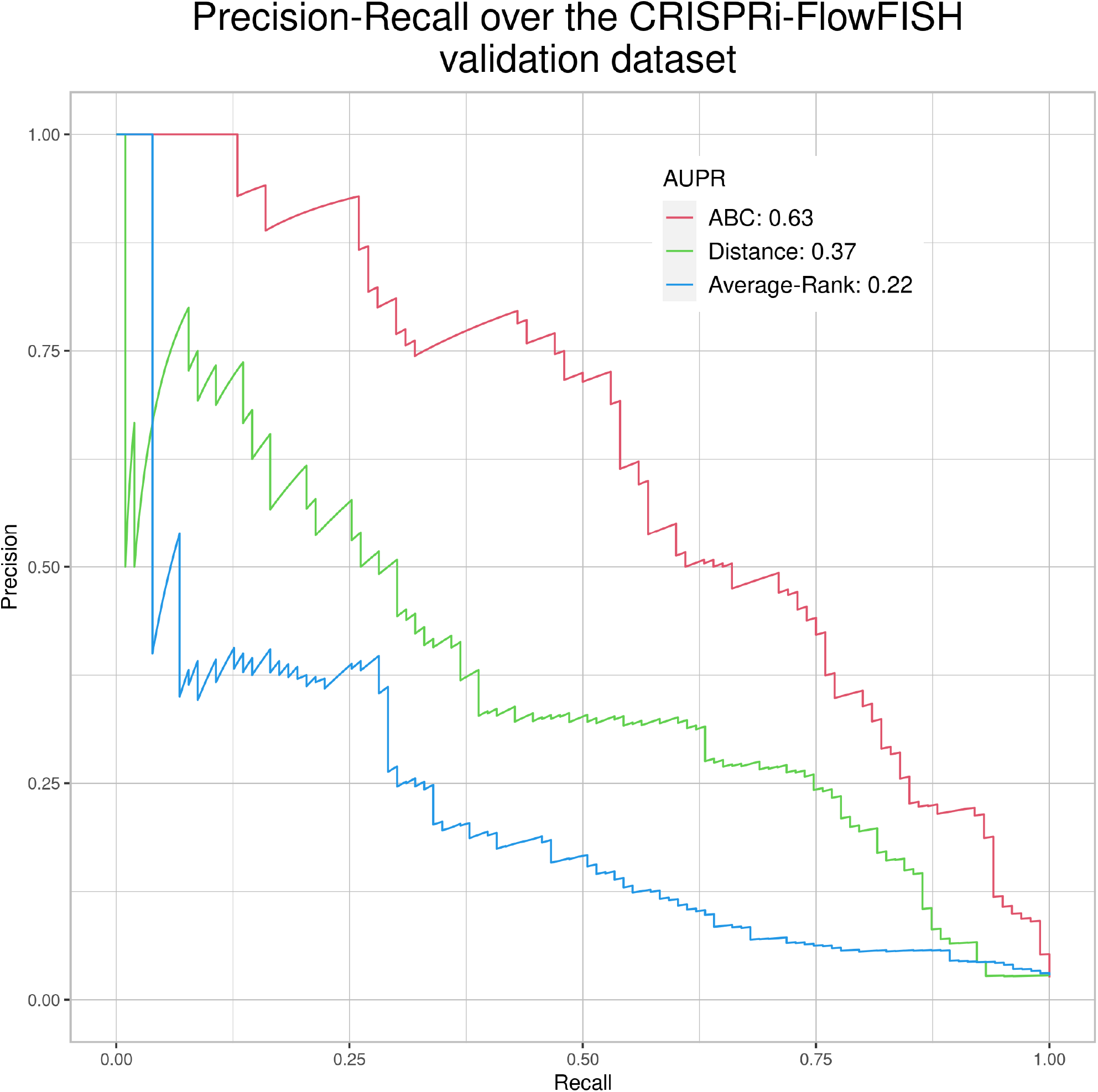
Precision-Recall curves for the `ABC model`, the `distance` and the `Average-Rank` methods on the CRISPRi-FlowFISH (`CRiFF`) evaluation set. AUPRs of each method are indicated on the top right end corner of the plot.

First, the results of the two evaluation papers ([12, 22]) were reproduced here: the curves and AUPRs of the `Average-Rank` and the `distance` methods of Figure 4 correspond to the curves and AUPRs of the part of Figure S2 of [22] that correspond to the GM12878 cell line (all pairs, natural ratio), and the curves and AUPRs of the ABC model and the distance method of Figure 5 correspond to the curves and AUPRs of Figure 3a of [12]. Altogether, these positive controls confirmed the validity of the pipeline we implemented.

More precisely, Figure 4 shows quite low AUPR values for the three methods on the six `BENGI` datasets, and that the `Average-Rank` method performs best, before the `distance` method and then the `ABC model`. On the other hand, Figure 5 shows larger AUPRs on the `CRiFF` set for the three methods and that the `ABC model` performs best (AUPR=0.63). before the `distance` method and then the `Average-Rank` method.

The bad performances (small precision values even for small recall values) of the `ABC model` on the `BENGI` sets could be due to the fact that the `BENGI` dataset’s underlying data (HiC, promoter capture HiC, ChiA-PET and RNA-seq) were not specifically designed to identify E/G relationships. Indeed there may be some E/G relationships that do not need spatial proximity or the presence of CTCF to operate [28]. Likewise the presence of an eQTL in a predicted enhancer does not necessarily imply the eQTL represents an E/G relationship.

The bad performance of the `Average-Rank` method (small precision values even for small recall values) on the `CRiFF` data is more difficult to explain as this technique should be quite exhaustive in identifying the enhancers of a given gene. However, the authors state in their paper that “CRISPRi might fail to discover certain regulatory elements, for example due to differential sensitivity to KRAB-mediated inhibition” ([12]). Why this would affect the `Average-Rank` method more than the `ABC model` still needs further investigation.

## Discussion

Looking for a good compromise between the two evaluations, at first glance the base-line `distance` method might appear as the best one, with the most stable results across evaluation sets. However we know this method does not work well in many cases ([18, 24, 25]). Therefore it is very difficult to decide on the best method based on the evaluation data we have here.

On the other hand, because it was specifically designed to identify E/G relationships for a selected set of genes, the `CRiFF` technique seems to be better suited to generate true E/G reference/evaluation data. Therefore if we really had to select an E/G relationship identification method, then we would choose the one that performs best on the `CRiFF` data, namely the `ABC model`. Another, more practical, reason to select the `ABC model` over the `Average-Rank` method is that a dedicated code has been made available to the community by their authors^4^, which is not the case for the `Average-Rank` method.

One of the drawbacks of the `CRiFF` technique however, is that it does not provide genome-wide results. In the study we have used here, it was run on thirty different genes located in five different genomic regions ranging from 1.1Mb to 4 Mb in size, and it can be wondered whether these genes are really representative of the whole genome. Another considerable bias could come from the use of the K562 cancer cell line which is the only cell line for which there was sufficient `CRiFF` data. Even if the authors have performed more CRISPRi-FlowFISH experiments since our study (283 true validated and 5756 false E/G in 11 cell types, [25]), this type of reference data is still not genome-wide and biased toward cancer cell lines.

Finally, altogether our results call for the generation of more complete and reliable E/G relationship reference/evaluation data, rather than for new more elaborate E/G relationship identification methods, such as the ones that are currently being developed. A more reliable genome-wide set of E/G would indeed allow to better evaluate the numerous already existing E/G relationship identification methods based on 1D data, in order to finally reach a consensus in this field, and be able to answer numerous questions related to cell function and disease.

## Material and Methods

### Pairwise chromatin accessibility correlation across cell types (Figure 2 and Table 2)

In order to represent the heuristic enhancer/gene identification methods, we chose the simplest one of them: the pairwise chromatin accessibility correlation across cell types, and represented it on Figure 2. Here are the actions that led to this figure: we first downloaded the ENCODE uniformly processed read alignments (`bam` files) of DNAse-seq data (single end) from 10 cell types: stomach, HepG2, K562, thymus, adrenal gland, small intestine, GM12878, IMR-90, heart and H1-hESC, with accession numbers provided in Table 2. These 10 `bam` files could easily be downloaded from the data search part of the ENCODE web site ^5^ We then called the chromatin accessibility peaks from the mapped reads in each cell type using macs2 version 2.1.1.20160309 and the following command:

**Table 2:**
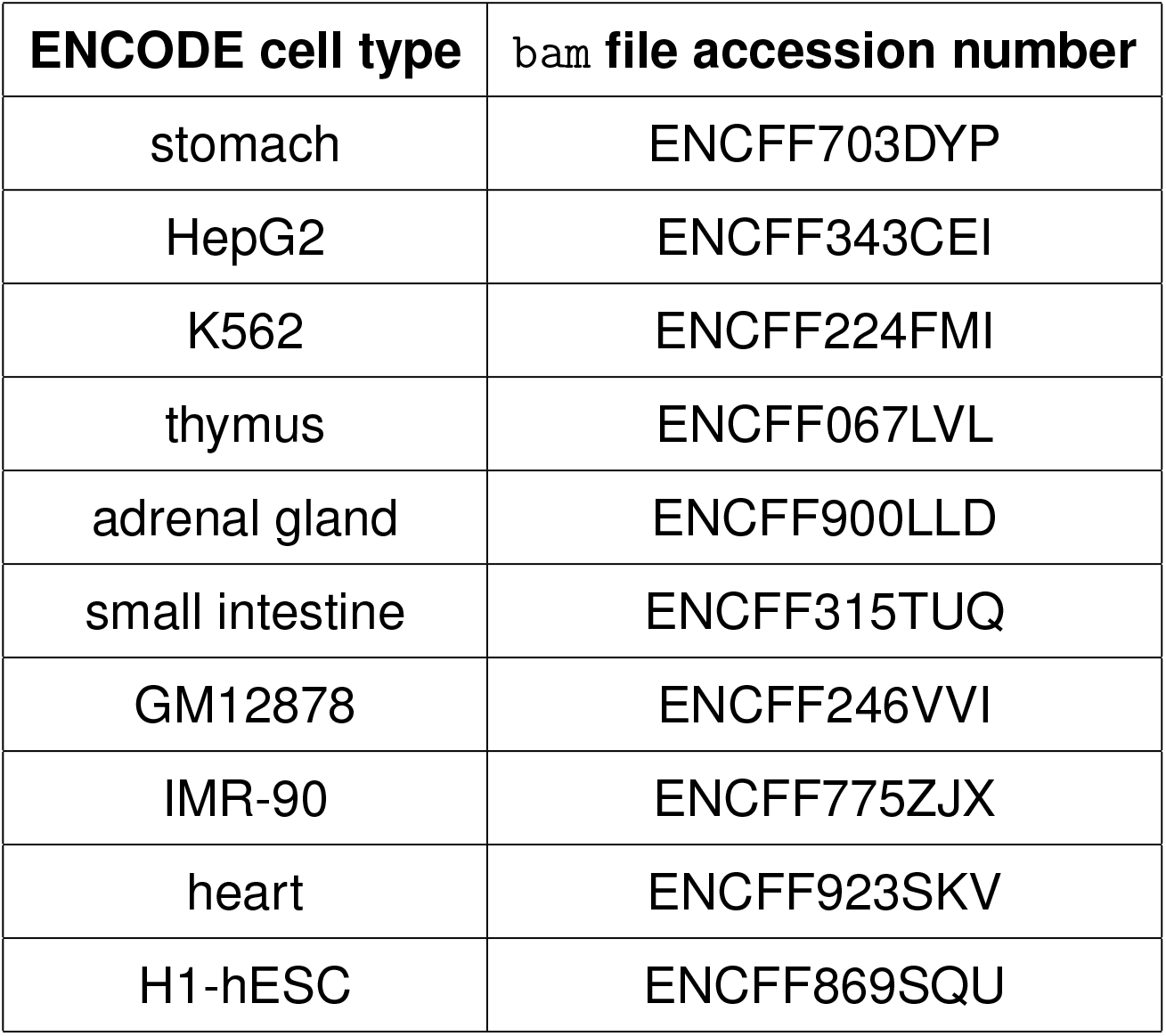
ENCODE cell types and accession numbers of associated DNA-seq alignment bam files.

~~~
macs2 callpeak --nomodel -f BAM -t $celltype.bam -n $celltype \
--keep-dup all --verbose 3 2> macs2.err > macs2.out
~~~

We obtained from about 60,000 (GM12878) to about 200,000 (IMR-90) peaks per cell type By concatenating, sorting and merging on the genome the peaks called in each cell type using `bedtools merge` version 2.29, we then obtained 473,766 consensus peaks across all cell types. We then quantified the chromatin accessibility of the 473,766 consensus peaks in each cell type by simply counting the number of mapped reads of each cell type overlapping each consensus peak using `bedtools intersect` version 2.29, and normalized the number of reads of each peak in each cell type by the total number of mapped reads in peaks for this cell type. Finally we computed the consensus peak pairwise Pearson correlation between the log10 of the normalized chromatin accessibility across the 10 cell types of these peaks for all pairs of peaks less distant than 500kb using a script that we wrote: `compute_correlations.py` from our github repository ^6^. We then only considered as E/G relationships the pairs of peaks with a correlation above 0.7 and for which one of the two peaks overlapped the most 5’ bp (TSS) of a Gencode v19^7^ gene (vertical green rectangles on Figure 2).

### Method evaluation (Table 1, Figures 4 and 5)

In addition to evaluating the `ABC model` on the `BENGI` sets and the `Average-Rank` method on the `CRiFF` set, and since Moore et al. provided the code of the `Average-Rank` method, and Fulco et al the one of the `ABC model`, we decided to try and reproduce the evaluation of the `Average-Rank` method on the `BENGI` sets and of the `ABC model` on the `CRiFF` set. We also used the code of the distance method provided by Moore et al, to evaluate the distance method on the `BENGI` and on the `CRiFF` sets, with some modifications. In total we evaluated three methods, the `ABC model`, the `Average-Rank` method and the distance method on two references sets, the `BENGI` and the `CRiFF` sets (Figures 4 and 5). It has to be noted that since the code to generate the Precision-Recall curves and the AUPRs was not provided in the papers, we generated our own R code to make these plots using existing R packages. The code used to perform all these analyses was stored in Jupyter notebooks that we provide below, together with additional details about these analyses.

### Method evaluation on the BENGI sets

The Moore et al’s code, reference data and annotation were first downloaded from the `BENGI` github repository ^8^. More precisely the `Scripts` directory included, on the one hand the scripts to make the `BENGI` sets, and on the other hand the scripts to run the evaluation of the methods on a given `BENGI` set (note that other cell types than GM12878 were provided). It is important to bear in mind that the script corresponding to a method was not a generic script allowing to retrieve all the E/G relationships called by this method in a particular cell type, but rather only produces evaluation data of this method on a given `BENGI` set, i.e. attaches to each true and false E/G of a `BENGI` set, the score of the method’s associated prediction (to be used to draw the Precision-Recall curves and compute the AUPRs). Since we could not run any of the scripts from Moore et al without modifying them, sometimes quite deeply, we suspect these scripts were provided to give a general idea of the underlying analyses (and satisfy the reviewers) rather than to be used as such. No mention of program versions were provided neither, which again hampers reproducibility.

### Evaluating the distance method on the BENGI sets

To evaluate the distance method on the `BENGI` sets, we used a slightly modified version of the `Run-Distance-Method.sh` script provided by Moore et al. This script takes as input a string defining the `BENGI` set (`celltype.settype`, for instance `GM12878.CHiC`), the version of the `BENGI` set (here `v3`), the mode (here `normal`), the expression threshold (here `0.2` but this parameter is not used in normal mode) and the output path. In normal mode, this script calls the `rank.distance.py` script on the set of human TSSs, the set of all `cCREs` and the `BENGI` set. It then outputs a 2 column file including for each E/G of the `BENGI` set on a row, 1 or 0 according to whether this E/G is true or false according to the `BENGI` set and the score provided by the `distance` method which is defined as the inverse of the smallest distance between a TSS of G and the enhancer E. Our modification consisted in adding two additional columns to this tabulated file, one for the enhancer id and one for the gene id, this for an easier downstream fusion with the evaluation result of the `Sheffield` method. For this purpose we also had to modify the `Run-Distance-Method.sh` script so that it sorts the 4 column tabulated file provided by the python script according to the enhancer id and the gene id. After running the evaluation script we plotted the Precision-Recall curves using existing R packages. The following Jupyter notebook provides all the necessary information for evaluating the `distance` method on the `BENGI` sets ^9^.

### Evaluating the Average-Rank method on the BENGI sets

To evaluate the `Average-Rank` method on the `BENGI` sets, we first had to run the `Sheffield` method (correlation between open chromatin at E and expression level at G) on each `BENGI` set.

For this we first downloaded the DNAse Hypersensitivity (DHS) peaks with their chromatin accessibility in 112 cell types (`dhs112_v3.bed` file) and the genes with their expression levels in the same 112 cell types (`exp112.bed` file) from the web ^10^ and as indicated in page 14 of [22]. We then ran the `Run-Sheffield.sh` script that evaluates the `Sheffield` method on a given `BENGI` set. This script takes as input a string defining the `BENGI` set, the version of the `BENGI` set and the output path. It then makes the set of enhancers of the `BENGI` set in `bed` format, the enhancer matrix with these enhancers in rows and their chromatin accessibility in the 112 cell types in columns, the genes of the `BENGI` set in `bed` format, the matrix of these genes in rows with their expression levels in the 112 cell types in columns, and then calls the `sheffield.correlation.py` script. This script takes as input a matrix of gene expression in the 112 cell types, the gene file in bed format, the enhancer matrix, a gene summary file, the `BENGI` set and the cell type. It then outputs a 6 column file including for each E/G of the `BENGI` set on a row, 1 or 0 according to whether this E/G is true or false in the `BENGI` set, the Pearson correlation between the chromatin accessibility at E and the expression level at G across the 112 cell types, the P-value, the Z-score, the enhancer id and the gene id.

In fact we had to modify the `Run-Sheffield.sh` script in order to 1) replace spaces by tabs in the DHS peak chromatin accessibility matrix, 2) have a `gene id <tab> gene name` file instead of a gene `bed` file, 3) replace spaces by tabs in the gene expression level matrix, 4) do without the gene summary file (that we could not find anywhere) and 5) sort the tabulated output according to enhancer id and gene id. Accordingly we also had to modify the `sheffield.correlation.py` script: 1) we added an `all equal` function to prevent divisions by zero in the `Calculate_Correlation` function and 2) we added code to create the gene summary file that the script was supposed to take as input. The complete process to run the `Sheffield` method on the `BENGI` set can be found on this page ^11^

Finally we ran the `Run-Average-Rank.sh` script that evaluates the `Average-Rank` method on a `BENGI` set. This script takes as input a string defining the `BENGI` set and the version of the `BENGI` set, and outputs a 7 column tabulated file including for each E/G of the `BENGI` set, 1 or 0 according to whether this E/G is true or false in the `BENGI` set, the average rank score, the distance score, the correlation score, the distance rank, the correlation rank and the average rank between the distance and the correlation. Here we also had to modify the bash script, first because the `distance` script now outputs a 4 instead of a 2 column file, but second and more importantly because we spotted a mistake in the calculation of the correlation rank, where the inverse distance is taken instead of the correlation (we had to replace $2 by $3 in the following portion of the code

~~~
sort -k3,3gr | awk ‘BEGIN{x=0;r=0}
{if (x != $2) r +=1; print $0 “\t” r “\t” ($NF+r)/2; x=$2}’
~~~

Once again we plotted the Precision-Recall curves using an R code of our own. The complete process to evaluate the `Average-Rank` method on the `BENGI` sets can be found here ^12^.

### Evaluating the ABC model on the BENGI sets

In order to evaluate the `ABC model` on the `BENGI` sets, we downloaded the `ABC model` code from its github repository ^13^. Although the complete process is not a pipeline but is rather made of several steps to run one after the other, the documentation was so pedagogic and complete that we had no particular issue running the `ABC model` on GM12878 data. We also found the tools and associated version to use. In case of doubt we wrote to the authors who always gave us a fast and precise answer. Nonetheless and for the sake of reproducibility the complete process is detailed in this notebook ^14^.

### Method evaluation on the CRISPRi-FlowFISH (CRiFF) set

To obtain the `CRiFF` reference set we first downloaded Table S6a from Fulco et al [12] as a tsv file, and then obtained the 109 positive and the 3754 negative E/G relationships performing the following filters (the negatives are indeed defined as either not significant or not associated to a decrease in gene expression):

~~~
awk ‘NR==2{gsub(/\ /,”.”,$0); header=$0; OFS=“\t”;
print header > “109.fulco.positives.tsv”;
print header > “3754.fulco.negatives.tsv”}
NR>=3{
split($0,a,”\t”); if(a[6]!=“promoter”&&(a[10]==“TRUE”||a[17]>0.8))
{
if(a[10]==“TRUE”&&a[8]<0)
{
print > “109.fulco.positives.tsv”;
}
else
{
print > “3754.fulco.negatives.tsv”;
}}}’ tableS6a.tsv
~~~

In order to be able to use almost the same scripts as above for the `distance` and the `Average-Rank` methods, we first intersected the `CRiFF` set with the ENCODE `cCREs` provided and used by Moore et al. This process is described in the three notebooks below. We have to say that we only slightly modified the distance and `Average-Rank` method scripts used above for `BENGI` and GM12878 in order to run then on `CRiFF` and K562 (see notebooks below).

**Evaluating the** `distance` **method on the** `CRiFF` **set** The complete process for this evaluation is provided in the following notebook ^15^

**Evaluating the** `Average-Rank` **method on the** `CRiFF` **set** The complete process for this evaluation is provided in the following notebook ^16^

**Evaluating the** `ABC model` **on the** `CRiFF` **set** The complete process for this evaluation is provided in the following notebook ^17^

## Author Contributions

TH performed the entire method evaluation, under the supervision of SD. CM worked on the chromatin accessibility correlation method under the supervision of SF, TF and SD. HA, WLG, SF and TF provided critical views on the evaluation results. SD designed the study.

## Acknowledgments

We would like to thank the authors of ([12, 22]) for the detailed Material and Methods’ section of their papers, that allowed us to reproduce the curves and AUPRs of Figure 3a of [12], and the part of Figure S2 corresponding to the GM12878 cell line (all pairs, natural ratio) for [22]. We would also like to thank the authors of [12] for their reactivity answering the questions we had about their study.

## Footnotes

1 http://enhancer.lbl.gov/

2 https://github.com/broadinstitute/ABC-Enhancer-Gene-Prediction

3 https://github.com/weng-lab/BENGI

4 https://github.com/broadinstitute/ABC-Enhancer-Gene-Prediction

5 https://www.encodeproject.org/

6 https://github.com/sdjebali/EnhancerGene

7 https://www.gencodegenes.org/

8 https://github.com/weng-lab/BENGI

9 https://genoweb.toulouse.inra.fr/thoellinger/notes/notes_BENGI/distance_method/distance_evaluation_with_code.html

10 http://big.databio.org/papers/RED/supplement/

11 https://genoweb.toulouse.inra.fr/thoellinger/notes/notes_BENGI/dnase_expression_correlation/correlation_method_with_code.html

12 https://genoweb.toulouse.inra.fr/thoellinger/notes/notes_BENGI/avg_rank_method/avg_rank_method_with_code.html

13 https://github.com/broadinstitute/ABC-Enhancer-Gene-Prediction

14 https://genoweb.toulouse.inra.fr/thoellinger/notes/notes_ABC/BENGI/notebook_ABC_over_BENGIGM12878_from_ccRE_ELSs.html

15 https://genoweb.toulouse.inra.fr/thoellinger/notes/notes_BENGI/CRISPRi_FlowFISH/distance_method/distance_over_fulco_et_al_crispri.html

16 https://genoweb.toulouse.inra.fr/thoellinger/notes/notes_BENGI/CRISPRi_FlowFISH/avg_rank_method/avg_rank_method_with_code.html

17 http://genoweb.toulouse.inra.fr/thoellinger/notes/notes_ABC/K562/april_K562_56_genes/april_K562_56_genes.html

## References

[1] R. Andersson, C. Gebhard, I. Miguel-Escalada, I. Hoof, J. Bornholdt, M. Boyd, Y. Chen, X. Zhao, C. Schmidl, T. Suzuki, et al. An atlas of active enhancers across human cell types and tissues. Nature, 507(7493):455–461, 2014.

[2] D. Aran, S. Sabato, and A. Hellman. Dna methylation of distal regulatory sites characterizes dysregulation of cancer genes. Genome biology, 14(3):1–14, 2013.

[3] O. G. Bahcall. Gtex pilot quantifies eqtl variation across tissues and individuals. Nature Reviews Genetics, 16(7):375–375, 2015.

[4] P. S. Belokopytova, M. A. Nuriddinov, E. A. Mozheiko, D. Fishman, and V. Fishman. Quantitative prediction of enhancer–promoter interactions. Genome research, 30(1):72–84, 2020.

[5] S. Buchka, A. Hapfelmeier, P. P. Gardner, R. Wilson, and A.-L. Boulesteix. On the optimistic performance evaluation of newly introduced bioinformatic methods. Genome biology, 22(1):1–8, 2021.

[6] Q. Cao, C. Anyansi, X. Hu, L. Xu, L. Xiong, W. Tang, M. T. Mok, C. Cheng, X. Fan, M. Gerstein, et al. Reconstruction of enhancer–target networks in 935 samples of human primary cells, tissues and cell lines. Nature genetics, 49(10):1428–1436, 2017.

[7] G. Consortium, K. G. Ardlie, D. S. Deluca, A. V. Segrè, T. J. Sullivan, T. R. Young, E. T. Gelfand, C. A. Trowbridge, J. B. Maller, T. Tukiainen, et al. The genotype-tissue expression (gtex) pilot analysis: multitissue gene regulation in humans. Science, 348(6235):648–660, 2015.

[8] O. Corradin, A. Saiakhova, B. Akhtar-Zaidi, L. Myeroff, J. Willis, R. Cowper-Sal, M. Lupien, S. Markowitz, P. C. Scacheri, et al. Combinatorial effects of multiple enhancer variants in linkage disequilibrium dictate levels of gene expression to confer susceptibility to common traits. Genome research, 24(1):1–13, 2014.

[9] O. Corradin and P. C. Scacheri. Enhancer variants: evaluating functions in common disease. Genome medicine, 6(10):1–14, 2014.

[10] J. Ernst, P. Kheradpour, T. S. Mikkelsen, N. Shoresh, L. D. Ward, C. B. Epstein, X. Zhang, L. Wang, R. Issner, M. Coyne, et al. Mapping and analysis of chromatin state dynamics in nine human cell types. Nature, 473(7345):43–49, 2011.

[11] Y. Fan and B. Peng. Stackepi: identification of cell line-specific enhancer– promoter interactions based on stacking ensemble learning. BMC bioinformatics, 23(1):1–18, 2022.

[12] C. P. Fulco, J. Nasser, T. R. Jones, G. Munson, D. T. Bergman, V. Subramanian, S. R. Grossman, R. Anyoha, B. R. Doughty, T. A. Patwardhan, et al. Activity-by-contact model of enhancer–promoter regulation from thousands of crispr perturbations. Nature genetics, 51(12):1664–1669, 2019.

[13] T. A. Hait, D. Amar, R. Shamir, and R. Elkon. Focs: a novel method for analyzing enhancer and gene activity patterns infers an extensive enhancer–promoter map. Genome biology, 19(1):1–14, 2018.

[14] B. He, C. Chen, L. Teng, and K. Tan. Global view of enhancer–promoter interactome in human cells. Proceedings of the National Academy of Sciences, 111(21):E2191–E2199, 2014.

[15] Z. Hong, X. Zeng, L. Wei, and X. Liu. Identifying enhancer–promoter interactions with neural network based on pre-trained dna vectors and attention mechanism. Bioinformatics, 36(4):1037–1043, 2020.

[16] I. Jung, A. Schmitt, Y. Diao, A. J. Lee, T. Liu, D. Yang, C. Tan, J. Eom, M. Chan, S. Chee, et al. A compendium of promoter-centered long-range chromatin interactions in the human genome. Nature genetics, 51(10):1442–1449, 2019.

[17] N. Kerimov, J. D. Hayhurst, K. Peikova, J. R. Manning, P. Walter, L. Kolberg, M. Samoviča, M. P. Sakthivel, I. Kuzmin, S. J. Trevanion, et al. A compendium of uniformly processed human gene expression and splicing quantitative trait loci. Nature genetics, 53(9):1290–1299, 2021.

[18] I. Krivega and A. Dean. Enhancer and promoter interactions—long distance calls. Current opinion in genetics & development, 22(2):79–85, 2012.

[19] T. Lappalainen, M. Sammeth, M. R. Friedländer, P. A. t Hoen, J. Monlong, M. A. Rivas, M. Gonzalez-Porta, N. Kurbatova, T. Griebel, P. G. Ferreira, et al. Transcriptome and genome sequencing uncovers functional variation in humans. Nature, 501(7468):506–511, 2013.

[20] W. Li, W. H. Wong, and R. Jiang. Deeptact: predicting 3d chromatin contacts via bootstrapping deep learning. Nucleic acids research, 47(10):e60–e60, 2019.

[21] B. Mifsud, F. Tavares-Cadete, A. N. Young, R. Sugar, S. Schoenfelder, L. Ferreira, S. W. Wingett, S. Andrews, W. Grey, P. A. Ewels, et al. Mapping long-range promoter contacts in human cells with high-resolution capture hi-c. Nature genetics, 47(6):598–606, 2015.

[22] J. E. Moore, H. E. Pratt, M. J. Purcaro, and Z. Weng. A curated benchmark of enhancer-gene interactions for evaluating enhancer-target gene prediction methods. Genome biology, 21(1):1–16, 2020.

[23] J. E. Moore, M. J. Purcaro, H. E. Pratt, C. B. Epstein, N. Shoresh, J. Adrian, T. Kawli, C. A. Davis, A. Dobin, R. Kaul, et al. Expanded encyclopaedias of dna elements in the human and mouse genomes. Nature, 583(7818):699–710, 2020.

[24] M. R. Mumbach, A. T. Satpathy, E. A. Boyle, C. Dai, B. G. Gowen, S. W. Cho, M. L. Nguyen, A. J. Rubin, J. M. Granja, K. R. Kazane, et al. Enhancer connectome in primary human cells identifies target genes of disease-associated dna elements. Nature genetics, 49(11):1602–1612, 2017.

[25] J. Nasser, D. T. Bergman, C. P. Fulco, P. Guckelberger, B. R. Doughty, T. A. Patwardhan, T. R. Jones, T. H. Nguyen, J. C. Ulirsch, F. Lekschas, et al. Genome-wide enhancer maps link risk variants to disease genes. Nature, 593(7858):238–243, 2021.

[26] L. A. Pennacchio, W. Bickmore, A. Dean, M. A. Nobrega, and G. Bejerano. Enhancers: five essential questions. Nature Reviews Genetics, 14(4):288–295, 2013.

[27] S. S. Rao, M. H. Huntley, N. C. Durand, E. K. Stamenova, I. D. Bochkov, J. T. Robinson, A. L. Sanborn, I. Machol, A. D. Omer, E. S. Lander, et al. A 3d map of the human genome at kilobase resolution reveals principles of chromatin looping. Cell, 159(7):1665–1680, 2014.

[28] H. Ray-Jones and M. Spivakov. Transcriptional enhancers and their communication with gene promoters. Cellular and Molecular Life Sciences, 78(19):6453–6485, 2021.

[29] C. Rödelsperger, G. Guo, M. Kolanczyk, A. Pletschacher, S. Köhler, S. Bauer, M. H. Schulz, and P. N. Robinson. Integrative analysis of genomic, functional and protein interaction data predicts long-range enhancer-target gene interactions. Nucleic acids research, 39(7):2492–2502, 2011.

[30] S. Roy, A. F. Siahpirani, D. Chasman, S. Knaack, F. Ay, R. Stewart, M. Wilson, and R. Sridharan. A predictive modeling approach for cell line-specific long-range regulatory interactions. Nucleic acids research, 43(18):8694–8712, 2015.

[31] S. Schoenfelder and P. Fraser. Long-range enhancer–promoter contacts in gene expression control. Nature Reviews Genetics, 20(8):437–455, 2019.

[32] N. C. Sheffield, R. E. Thurman, L. Song, A. Safi, J. A. Stamatoyannopoulos, B. Lenhard, G. E. Crawford, and T. S. Furey. Patterns of regulatory activity across diverse human cell types predict tissue identity, transcription factor binding, and long-range interactions. Genome research, 23(5):777–788, 2013.

[33] Y. Shen, F. Yue, D. F. McCleary, Z. Ye, L. Edsall, S. Kuan, U. Wagner, J. Dixon, L. Lee, V. V. Lobanenkov, et al. A map of the cis-regulatory sequences in the mouse genome. Nature, 488(7409):116–120, 2012.

[34] Z. Tang, O. J. Luo, X. Li, M. Zheng, J. J. Zhu, P. Szalaj, P. Trzaskoma, A. Magalska, J. Wlodarczyk, B. Ruszczycki, et al. Ctcf-mediated human 3d genome architecture reveals chromatin topology for transcription. Cell, 163(7):1611–1627, 2015.

[35] R. E. Thurman, E. Rynes, R. Humbert, J. Vierstra, M. T. Maurano, E. Haugen, N. C. Sheffield, A. B. Stergachis, H. Wang, B. Vernot, et al. The accessible chromatin landscape of the human genome. Nature, 489(7414):75–82, 2012.

[36] S. Whalen, R. M. Truty, and K. S. Pollard. Enhancer–promoter interactions are encoded by complex genomic signatures on looping chromatin. Nature genetics, 48(5):488–496, 2016.

[37] Y. Yang, R. Zhang, S. Singh, and J. Ma. Exploiting sequence-based features for predicting enhancer–promoter interactions. Bioinformatics, 33(14):i252–i260, 2017.

[38] L. Yao, H. Shen, P. W. Laird, P. J. Farnham, and B. P. Berman. Inferring regulatory element landscapes and transcription factor networks from cancer methylomes. Genome biology, 16(1):1–21, 2015.

[39] G. Zhang, J. Shi, S. Zhu, Y. Lan, L. Xu, H. Yuan, G. Liao, X. Liu, Y. Zhang, Y. Xiao, et al. Diseaseenhancer: a resource of human disease-associated enhancer catalog. Nucleic acids research, 46(D1):D78–D84, 2018.

